# Two components of rapid temporal recalibration: a stimulus-driven shift and a decisional carryover dissociate across the senses

**DOI:** 10.64898/2026.09.29.755365

**Authors:** John Cass, Erik Van der Burg

## Abstract

Multisensory synchrony judgments depend on recent history: the point of subjective simultaneity (PSS) shifts toward recently experienced timing relationships. This shift, known as rapid temporal recalibration, is almost always interpreted as perceptual, a change in when events appear simultaneous. Yet serial dependence in such judgments can arise from two mechanistically distinct sources that a raw PSS shift cannot separate: a genuine perceptual realignment, which slides the psychometric function sideways along the multisensory asynchrony axis (an odd effect), and a decisional carryover, a bias to repeat the previous verdict, which lifts the function vertically without moving the PSS (an even effect). We re-analysed eight published datasets investigating rapid recalibration spanning audiovisual, audiotactile, and visuotactile pairings and audiovisual speech, decomposing serial dependence into these two components within a single mixed-effects model. The components dissociated by modality: perceptual recalibration was robust for audiovisual pairs but weak for audiotactile and absent for visuotactile pairings, whereas the decisional carryover was largest for the tactile pairings and weak for audiovisual pairings, so that the two traded off across the senses. In audiovisual speech, the decisional carryover was gated by the spoken syllable but not by the talkers’ identity and did not persist beyond one trial, identifying it as a content-specific, one-step decision memory rather than a motor habit or a multi-trial store. Separating the perceptual and decisional contributions to a timing judgment thus re-draws the map of temporal recalibration and revises several conclusions drawn from the undecomposed PSS.

## Introduction

To perceive events as unified, the brain must contend with the fact that the signals composing them arrive at the sensory receptors, and at the cortex, at different times Hirsh and Sherrick (1961), (Poppel et al., 1990). For example, a flash and a click from a common source may reach an observer at different times because light and sound travel at different speeds, while additional temporal offsets arise from differences in neural processing across sensory modalities (Keetels and Vroomen, 2012). Despite these differences, signals originating from the same event are typically experienced as simultaneous. One way the system achieves this is by continually recalibrating its estimate of cross-modal simultaneity, shifting the point of subjective simultaneity (PSS) toward recently experienced timing relationships (Fujisaki et al., 2004, Vroomen et al., 2004). Such recalibration has been documented over two apparently distinct timescales: a slow, cumulative adaptation that develops over minutes of exposure and outlasts it as an aftereffect (Fujisaki et al., 2004, Vroomen et al., 2004), and a rapid, trial-by-trial adjustment in which a single preceding stimulus shifts judgments on the very next trial (Van der Burg et al., 2015b, Van der Burg et al., 2018, Van der Burg et al., 2013).

These effects are almost universally measured as a displacement of the PSS. It is tempting to read such a displacement as perceptual, a change in when (at what physical asynchrony) the observer experiences two events as temporally coincident. However, a shift in the PSS does not, by itself, establish such a perceptual change. As Yarrow and colleagues have argued, a PSS displacement can instead reflect a shift in the decision criteria that define which asynchronies are judged synchronous, with no change in perceptual timing at all (Yarrow et al., 2011, Yarrow et al., 2015). A perceptual realignment and a re-drawn synchrony boundary both slide the psychometric function along the SOA axis, so they cannot be distinguished from the PSS on its own. This is one sense in which recalibration may be “decisional.”

Our decomposition is aimed at a second, logically distinct source of decisional influence that PSS-based analyses miss entirely. Beyond any shift in where the function peaks, serial dependence can express itself as an overall bias to repeat the previous judgment, increasing or decreasing the probability of a “synchronous” response irrespective of the current SOA. This repetition bias is decisional in a different sense from the criterion shifts described by Yarrow and colleagues. It reflects a carryover of the prior verdict rather than a re-drawing of the synchrony window, and it does not move the PSS. Importantly, these two sources are formally separable. A shift in the psychometric function relating “synchronous” responses to SOA appears as an interaction between the previous trial and current SOA (an odd effect, antisymmetric about the PSS), whereas a repetition bias appears as a main effect of the previous trial (an even effect, that is symmetric around the PSS). Standard analyses collapse across this odd/even distinction and thereby leave unresolved precisely how recent history shapes subsequent judgments.

The distinction is consequential. If part of what has been called rapid recalibration is a repetition bias rather than a shift in perceived timing, then its computational interpretation (what is updated, over what representation, and whether it reflects perception at all) changes accordingly, as does its relationship to the slow adaptation with which it is often grouped. Whether the recalibration itself reflects perceptual alignment or a change in decision criteria in Yarrow’s sense remains debated (Roseboom, 2019, Yarrow et al., 2011, Van der Burg et al., 2013, Van der Burg et al., 2018, Keane et al., 2019), and the odd/even decomposition does not resolve this question. What it does do, for the first time systematically, is isolate the repetition-bias component and separate it from the SOA-dependent shift across modalities and paradigms. We therefore re-analyse a family of temporal-recalibration datasets spanning simple audiovisual, audiotactile, and visuotactile stimulus pairings, as well as more complex audiovisual speech (Van der Burg et al., 2013, Van der Burg and Goodbourn, 2015, Van der Burg et al., 2015b, Van der Burg et al., 2015a), under a single decomposition that estimates the odd (shift) and even (repetition-bias) components within one model. The resulting pattern reveals a striking dissociation: the SOA-dependent shift is robust for audiovisual timing, weak for audiotactile, and absent for visuotactile, whereas repetition-bias carryover is largest precisely where the shift is weakest.

A decomposition is only as useful as the psychological reality of its parts, and the two components make divergent predictions about content-specificity. A low-level perceptual realignment should be tied to stimulus timing and largely indifferent to what was presented. A decisional carryover, being a bias about the judgment itself, could instead be gated by whether the current stimulus matches the preceding one. The audiovisual speech experiments of Van der Burg and Goodbourn (2015) are potentially informative here, because across trials they varied two things that are logically separable: *what* was said, the syllable that carries the very timing relationship being judged, and *who* said it, the identity of the talker, which is incidental to that relationship. This maps onto a distinction between the content, the message on which the synchrony judgment is based, and the messenger who conveys it. If the even effect is a genuinely decisional carryover, a verdict about synchrony carried forward, then it should be gated by the message but not the messenger: persisting when the syllable repeats, diminishing or disappearing when the syllable changes, yet surviving a change of talker.

We therefore, (i) decompose serial dependence in temporal judgments into perceptual and decisional components across eight datasets and three modality pairings; (ii) characterise the content-specificity of the decisional component, showing that it is gated by phonetic content (syllable) but invariant to talker identity, and that it operates one trial back; (iii) examine how the two components trade off across modalities and experimental paradigms.

## Method

### Datasets and participants

We reanalysed eight publicly or previously reported temporal-recalibration datasets from a common experimental lineage, spanning three modality pairings and audiovisual speech. In every study observers made a binary simultaneity judgment (“synchronous” versus “asynchronous”) on a multisensory stimulus presented at one of a fixed set of stimulus onset asynchronies (SOAs). Although recent stimulus history was not experimentally manipulated, the sequential structure of the data allowed each judgment on trial *t* to be analysed as a function of the stimulus configuration on the preceding trial (*t* − 1), and, where relevant, on earlier trials. In particular, trials were classified according to the modality order on the preceding trial, allowing us to quantify how recent temporal experience influenced the probability of judging the current stimulus as synchronous.

The datasets were: an audiovisual rapid-recalibration study (Van der Burg et al., 2013), labelled here as [AV-2013]; audiovisual, audiotactile, and visuotactile studies from a single cross-modal report (Van der Burg et al., 2015b) [labelled here as AV-2015b, AT-2015b, VT-2015b]; an audiovisual study (Van der Burg et al., 2015a) involving prolonged and trial-by-trial recalibration [labelled here as AV-2015a] (only the trial-by-trial data are analysed here); and three audiovisual speech experiments in which successive trials could repeat or switch the spoken syllable and the talker (Van der Burg and Goodbourn, 2015) [labelled AV-speech E1 to E3]. Participant numbers, SOA levels, and trial counts are summarised in Table 1. The exclusion criteria applied to each dataset are detailed below and in the Table 1 caption.

**Table 1.** Summary of the eight datasets included in the reanalysis. For each dataset, the table reports the source study, modality pairing, number of participants retained after application of the original study exclusion criteria, number and signed range of stimulus onset asynchrony (SOA) levels, and number of trials included in the present analyses after removal of the first trial of each block and trials with missing history information. Positive SOA values denote the visual modality leading for audiovisual and visuotactile pairings, and the tactile modality leading for audiotactile pairings. For the 2015b datasets, trials whose preceding trial had an SOA of 0 ms are additionally excluded (see Methods).

| Dataset | Source | Modality |  | N | SOA levels (ms) | Trials |
| --- | --- | --- | --- | --- | --- | --- |
|  |  | pairing |  |  |  |  |

| Dataset | Source | Modality |  | SOA levels (ms) | Trials |
| --- | --- | --- | --- | --- | --- |
|  |  | pairing | N |  |  |
| **AV-2013** | Van der Burg, Alais & Cass (2013) | Audiovisual | 15 | 9 (−512 to 512) | 17,556 |
| **AV-2015b** | Van der Burg, Orchard-Mills & Alais (2015) | Audiovisual | 18 | 7 (−424 to 424) | 9,626 |
| **AT-2015b** | Van der Burg, Orchard-Mills & Alais (2015) | Audiotactile | 18 | 7 (−424 to 424) | 9,628 |
| **VT-2015b** | Van der Burg, Orchard-Mills & Alais (2015) | Visuotactile | 18 | 7 (−424 to 424) | 9,629 |
| **AV-2015a** | Van der Burg, Alais & Cass (2015) | Audiovisual | 10 | 9 (−512 to 512) | 31,040 |
| **AV-speech E1** | Van der Burg & Goodbourn (2015) | Audiovisual (speech) | 10 | 11 (−800 to 800) | 7,600 |
| **AV-speech E2** | Van der Burg & Goodbourn (2015) | Audiovisual (speech) | 8 | 11 (−800 to 800) | 6,080 |
| **AV-speech E3** | Van der Burg & Goodbourn (2015) | Audiovisual (speech) | 8 | 11 (−800 to 800) | 14,440 |

For each dataset we applied the participant-level and trial-level exclusion criteria specified by its source study. For AV-2015a we retained the ten participants who completed the full multi- session protocol, matching the sample reported in the source study. Two further participants tested in a single session were not analysed. Across all datasets we additionally removed the first trial of every block, for which no previous-trial predictor is defined, and trials with missing responses or missing previous-trial values. For the three 2015b datasets we further excluded trials whose preceding trial had an SOA of 0 ms, for which no previous lead is defined, following the analysis convention of the source study; in the other datasets the previous SOA enters the models continuously and such trials were retained.

### Coding conventions and data preparation

Within each dataset, SOA was signed consistently so that positive values denote the visual (or, for tactile pairings, the second-named) modality leading and negative values the auditory (or first-named) modality leading. For each trial t, we derived three history predictors from the immediately preceding trial within the same block: the SOA on trial t-1, the sign of that SOA (i.e., the modality order on t-1), and the response on t-1 (“synchronous” versus “asynchronous”). For the audiovisual speech datasets we additionally coded whether the spoken syllable and the talker on trials t-1 and t-2 matched those on the current trial.

Continuous predictors entering the mixed-effects models were z-scored within dataset. Binary predictors (previous response, content match) were centred.

### Perceptual and decisional decomposition

The core decomposition rests on the parity of a history effect about the PSS. A change in the previous trial that shifts the psychometric function horizontally along the SOA axis appears as an interaction between the previous stimulus and the current SOA, an odd component that we label perceptual. A change that raises or lowers the overall probability of a “synchronous” response without moving the PSS appears as a main effect of the previous trial, an even component that we label decisional. These labels are geometric shorthand for where each effect acts on the psychometric function; whether the odd component is perceptual in a deeper mechanistic sense is taken up in the Discussion. We estimated both within a single generalised linear mixed model per dataset, fitted to trial-level binary responses with a logit link:

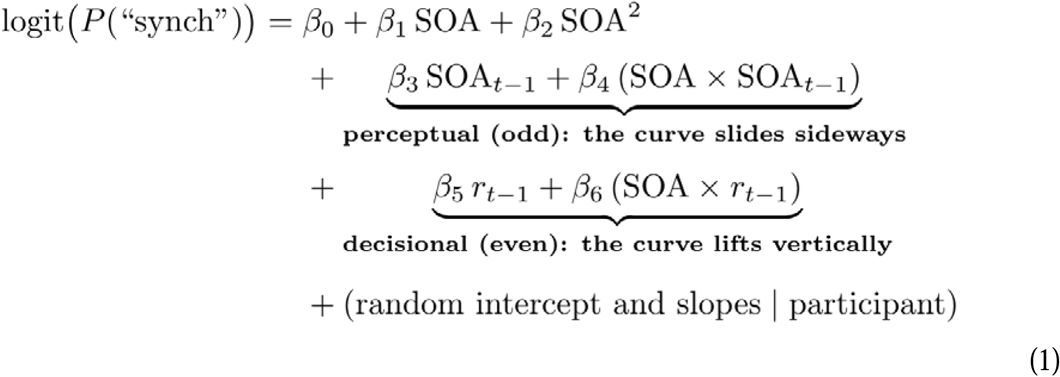

where the SOA×SOA_t-1_ interaction is the perceptual (odd) carryover and the t1_response main effect is the decisional (even) carryover. The quadratic SOA term captures the symmetric fall-off of the simultaneity function, and the SOA×t1_response term allows a history-dependent change in slope. Models were fitted in R using lme4 (glmer, bobyqa optimiser), with participant random intercepts and random slopes for the within-participant predictors; we report the perceptual and decisional coefficients with Wald tests, and, for the key contrasts, likelihood-ratio tests against the corresponding reduced model.

### Psychometric fits and effect magnitudes in interpretable units

To express each component in interpretable units and to visualise the split, we fitted Gaussian simultaneity functions of the form shown below:

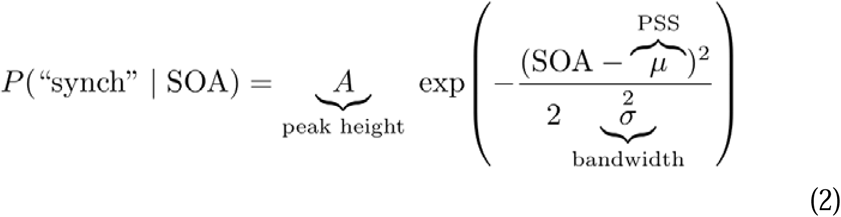

where μ is the PSS, σ the temporal bandwidth, and A the peak response probability. Parameters were estimated per participant by maximising the binomial likelihood of that participant’s counts at each SOA, rather than by unweighted least squares: likelihood-based fitting respects the [0,1] response range and weights each SOA level by its number of trials, avoiding the flattened fits least squares can produce when performance is near ceiling, that is, at SOA levels where almost every response is “synchronous”. Group-level display curves were fitted to the participant-averaged response proportions, weighted by trial count, so that they describe the same quantity as the plotted group means. Fits were computed separately for the two levels of the relevant history variable. For the perceptual component we split trials by the previous lead and took the perceptual effect as the horizontal separation of the two functions, ΔPSS = PSS(previous second- modality-led) − PSS(previous first-modality-led), in milliseconds. For the decisional component we split trials by the previous response and took the effect as the SOA-controlled vertical separation between the two conditions: at each SOA level we computed the difference in the proportion of “synchronous” responses following a previous “synchronous” versus a previous “asynchronous” response, and averaged these differences across SOA levels, weighting by trial count. SOA levels with fewer than five trials in either previous-response cell were omitted, so that no part of the average rests on a handful of trials. We used this model-free gap rather than the difference in fitted peak amplitude (Δpeak-P) because amplitude trades off against bandwidth in the Gaussian fits, which inflates the apparent decisional effect, particularly for the audiovisual datasets. Between-participant standard errors were computed across per-participant estimates.

### Content-specificity analyses (audiovisual speech stimuli)

For the speech datasets we tested whether the decisional carryover depends on the repetition of stimulus content. Within a logistic model of the current response including SOA and SOA², we added the previous response, an indicator of whether the previous trial matched the current trial on the relevant feature (syllable in Exp.1, talker in Exp.2, congruency in Exp.3), and their interaction; a reliable previous-response-by-match interaction indicates content-gated carryover.

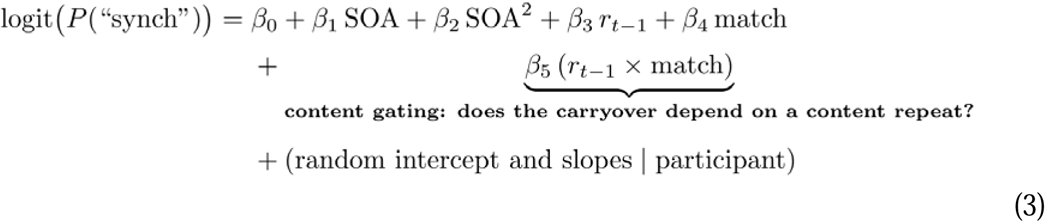

To ask whether any gated carryover reaches across an intervening non-matching trial, we extended the model to include the response two trials back and its interaction with two-back content match, and, as a targeted test, estimated the two-back carryover separately among trials on which the one-back trial was a content switch. All content analyses controlled for current SOA and were fitted on the trial-level responses. Trials without a defined two-back predictor (the first two trials of each block) were excluded from the two-back analyses.

### Cross-dataset relationship between components

To examine how the two components covary, we estimated, for each participant in each dataset, the perceptual and decisional coefficients from a penalised (Firth) logistic fit of the trial-level model to that participant’s data alone, and analysed the relationship across the resulting participant-by-dataset estimates.

### Software

Mixed-effects models were fitted in R 4.3.3 (Team, 2023) with lme4 1.1-35. Per-participant penalised (Firth, 1993) logistic fits used the brglm2 package 0.9.2. Psychometric fitting, content- gating, correlations, and figures were carried out in Python 3.11 (maximum-likelihood psychometric fits via SciPy’s optimiser; per-participant logistic models via statsmodels). All figures show individual participants at low opacity and group means with between-participant standard errors.

## Results

Every analysis begins from a simple idea about how the recent past can affect a simultaneity judgment. On each trial the probability of a “synchronous” response, plotted against the stimulus onset asynchrony (SOA), forms a bell-shaped function that peaks at the PSS. The previous trial can move this function in two geometrically distinct ways. It can slide it sideways along the SOA axis, changing the timing at which the two stimuli are most likely to be judged as simultaneous, which we call the perceptual (odd) component. Or it can lift it vertically, changing the overall probability of a “synchronous” response without moving the peak, which we call the decisional (even) component. These two changes are visible directly when the psychometric function is split by what happened on the previous trial.

Splitting each dataset’s function by the previous stimulus (whether the earlier trial had the first-named or second-named modality leading) separates the two curves horizontally (Fig. 1). A perceptual effect should appear as a horizontal separation between the two curves, reflecting a shift in the PSS (i.e., rapid temporal recalibration). ΔPSS is significant for all the audiovisual datasets, including the three speech experiments. For the audiotactile pair the shift is small (ΔPSS = +4 ms) and not significant in the participant-level test, although the mixed-effects model detects a small effect (p = .007; see below); for the visuotactile pair it is absent. Splitting instead by the previous response (whether the earlier verdict was “synchronous” or “asynchronous”) separates the curves vertically (Fig. 2). A decisional effect should instead appear as a vertical separation between the curves, reflecting a change in the overall probability of a “synchronous” response. Here the pattern reverses: the decisional effect is largest for the tactile pairings and close to zero for speech. The faint individual curves in both figures show that these are not artefacts of averaging; they hold within single participants.

**Figure 1.**
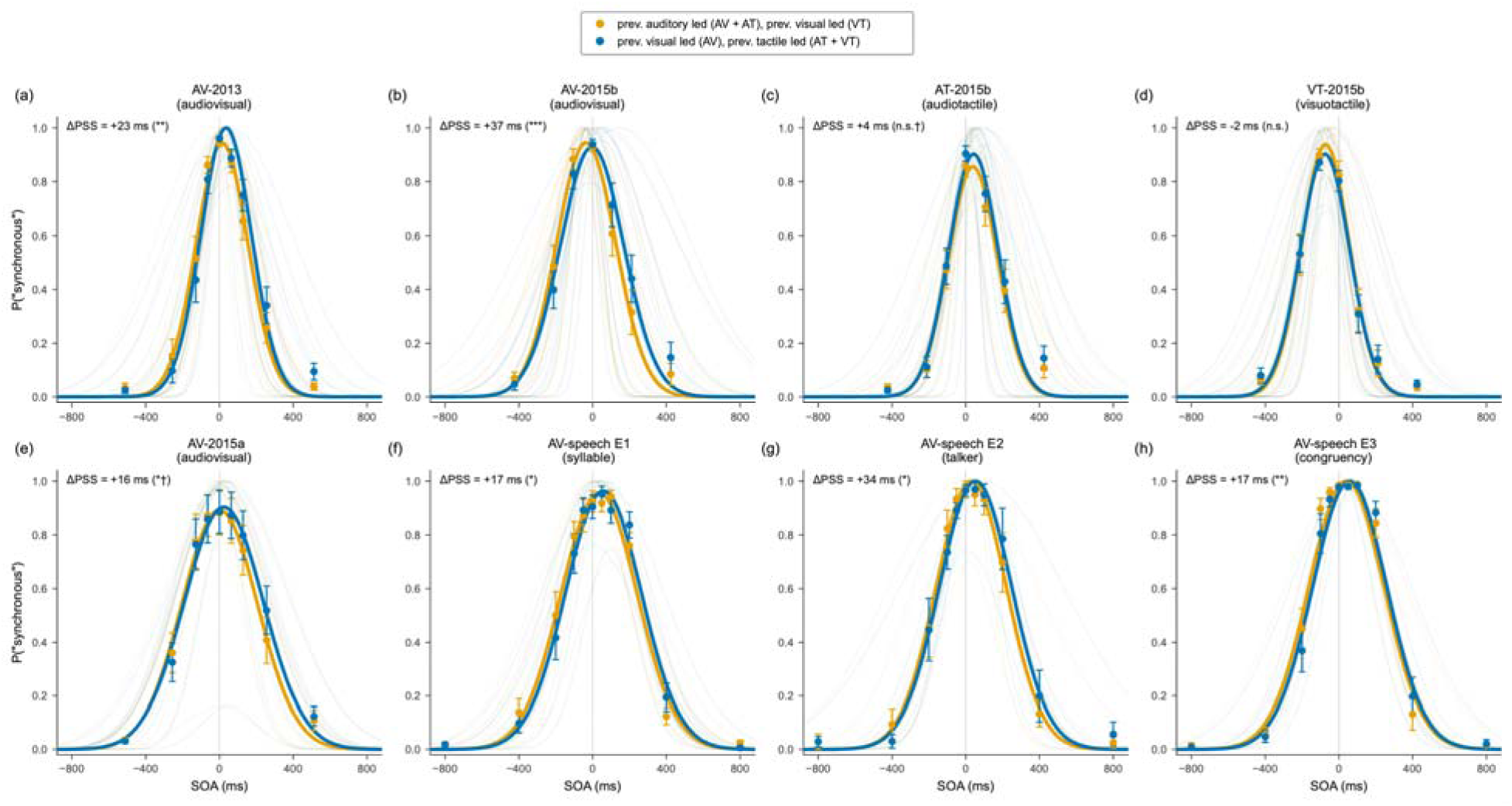
Simultaneity functions conditioned on the stimulus order of the immediately preceding trial, shown separately for all eight datasets. Here, the group mean probability of a “synchronous” response is shown as a function of the current-trial stimulus onset asynchrony (SOA) and leading modality on the previous trial. Orange symbols and curves correspond to trials preceded by an auditory-led stimulus in the audiovisual and audiotactile datasets, and by a visual-led stimulus in the visuotactile dataset; blue symbols and curves correspond to trials preceded by a visual-led stimulus in the audiovisual datasets, and by a tactile-led stimulus in the audiotactile an visuotactile datasets. Error bars represent the between-participant SEM, faint curves show participant-level fits, and bold curves show the corresponding group-level fits. The rapid temporal recalibration effect is expressed as the horizontal separation between the fitted functions, quantified as the difference in PSS (ΔPSS). Asterisks give a one-sample test across participants of the per-participant values summarised in each panel title (* *p* < .05, ** *p* < .01, *** *p* < .001); † denotes a dataset for which the maximal mixed-effects model yielded a different significance conclusion. All panels share a common SOA axis.

**Figure 2.**
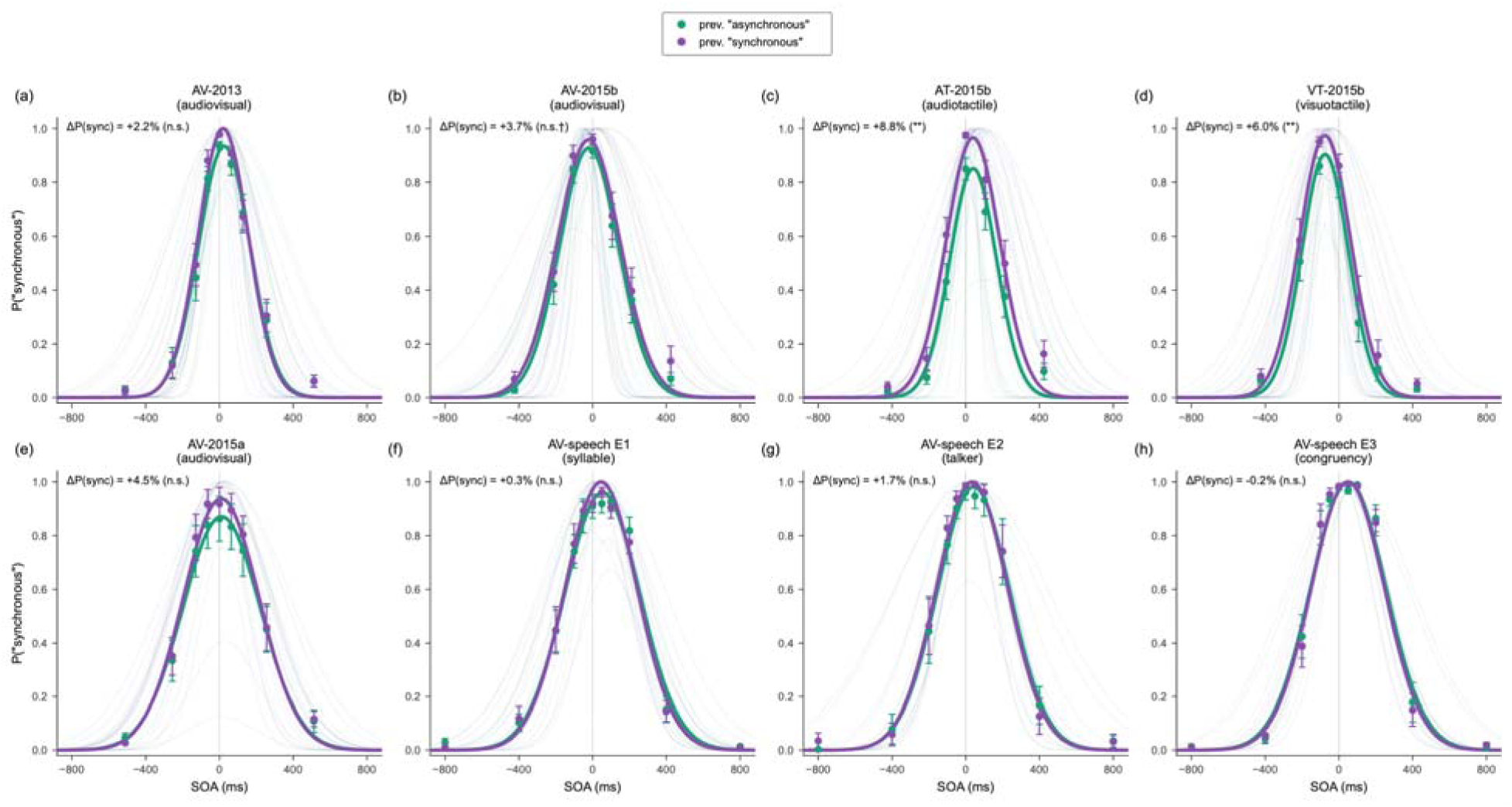
Simultaneity functions conditioned on the response given on the immediately preceding trial, show separately for all eight datasets. Here, the group mean probability of a “synchronous” response is shown as a function of the current trial stimulus onset asynchrony (SOA) and the preceding response. Green symbols and curves correspond to trials preceded by an “asynchronous” response; purple symbols and curves correspond to trials preceded by a “synchronous” response. Error bars represent the between-participant SEM, faint curves show participant-level fits, and bold curves show the corresponding group-level fits. The decisional carryover is quantified as the SOA-controlled difference in synchronous-response probability (ΔP(sync)), averaged across SOA levels. Asterisks give a one-sample test across participants of the per-participant values summarised in each panel title (* p < .05, ** p < .01, *** p < .001); † denotes a dataset in which the maximal mixed-effects model reaches the opposite verdict. All panels share a common SOA axis.

### Estimating the two components: the model

To estimate both components at once, and to test them properly, we fitted a single logistic mixed-effects model to the trial-by-trial responses of each dataset, shown as Eq. 1 in the Methods.

In Eq. 1 the perceptual carryover is the SOA × previous-stimulus interaction and the decisional carryover is the previous-response main effect. Every model included by-participant random slopes for these effects, so that each person is allowed their own version of each component and the test generalises to the population rather than to these particular trials. This specification matters: a pooled model that omits the random slopes treats every trial as independent and is strongly anti-conservative. Where we quote a significance level below, we confirmed it a second, independent way, by fitting each participant separately and testing the resulting estimates across the group.

To express the components in interpretable units we also fitted Gaussian simultaneity functions (Eq. 2), separately for each level of the relevant history variable, and read the perceptual effect as the horizontal shift of the peak (ΔPSS, in milliseconds) and the decisional effect as the SOA- controlled change in the probability of a “synchronous” response (ΔP(sync), in %).

### The shape of each component across the previous SOA

Plotting each component as a function of the previous trial’s SOA confirms the odd/even logic and reveals how the two components differ in form (Fig. 3). The perceptual component follows an odd, antisymmetric pattern: the previous SOA pushes the current PSS toward it, most strongly for intermediate SOAs. The marginal PSS, which combines perceptual and decisional influences, generally follows a similar pattern and closely tracks the perceptual component. By contrast, the decisional component follows an even, roughly symmetric function of the previous SOA, reflecting a bias that depends on the previous verdict rather than on the direction of the previous asynchrony. This dependence on previous SOA is expected because SOA strongly predicts th judgment made on the preceding trial: short asynchronies are more likely to be judged as synchronous, whereas large asynchronies are more likely to be judged as asynchronous. Thus, the decisional component depends on the previous verdict rather than on which modality led on the previous trial. The marginal PSS is close to the perceptual component in every dataset except the visuotactile one, where the two diverge precisely because there the recalibration is carried by the decisional rather than the perceptual component.

**Figure 3.**
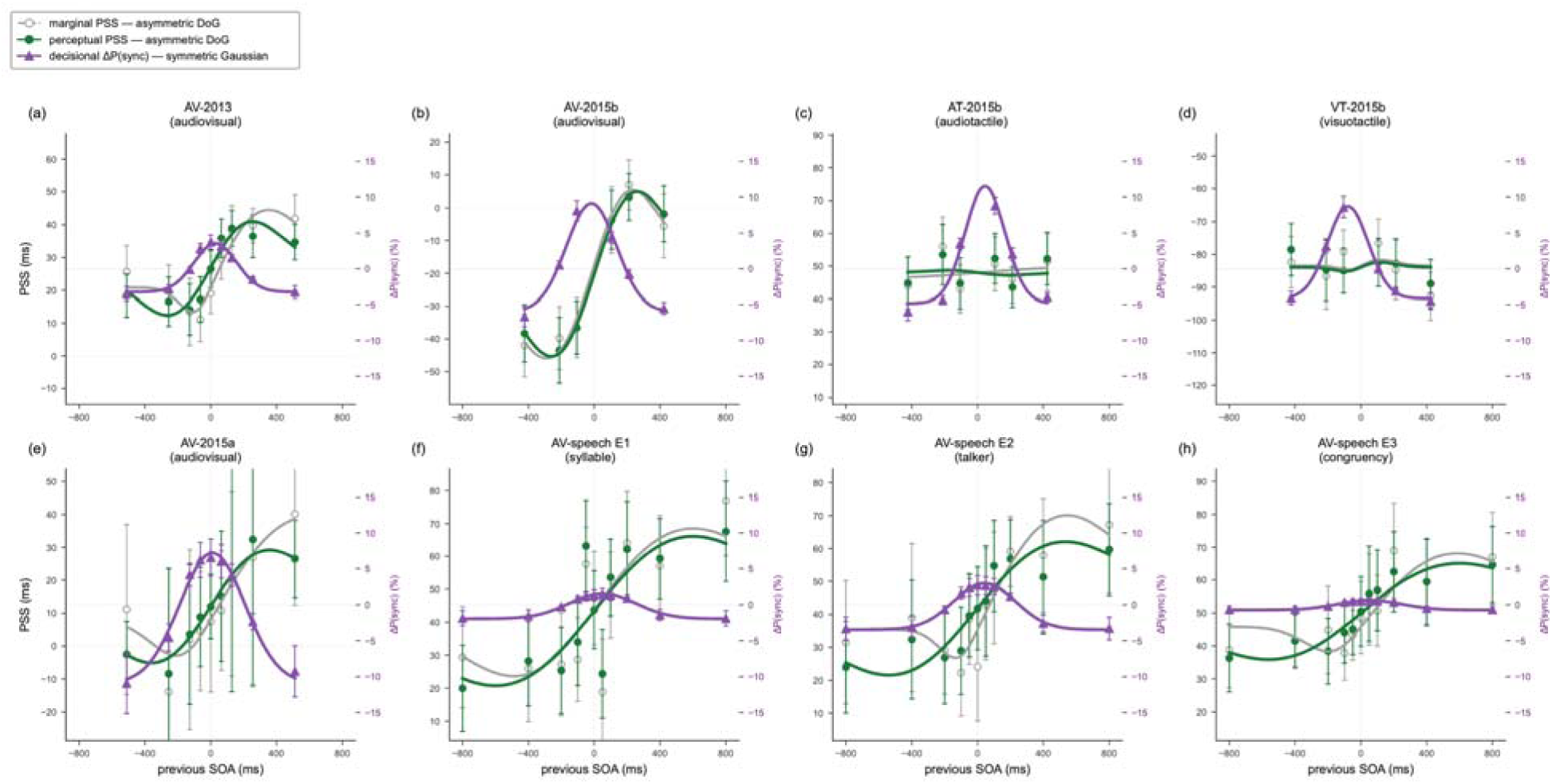
Dependence of the perceptual (ΔPSS) and decisional (ΔP(sync)) components on the stimulus onset asynchrony (SOA) of the preceding trial, shown separately for all eight datasets. The x-axis gives the previous trial’s SOA (ms). Grey symbols and curves show the marginal PSS estimated without separating perceptual and decisional contributions; green symbols and curves show the perceptual PSS component. Both are referenced to the left y-axis. Purple symbols and curves show the decisional carryover associated with the previous response, referenced to the right y-axis. Symbols show estimates at each previous-SOA level with error bars; solid curves show the corresponding fitted functions. All panels share a common previous-SOA axis; the left (PSS) axes share a common span and the right (ΔP(sync)) axes a common range, so magnitudes are comparable across panels.

### A double dissociation across the senses

Summarising both components across all eight datasets reveals a clean double dissociation (Fig. 4). The perceptual component was most consistent for audiovisual stimuli. The previous stimulus shifted the PSS by approximately 23 ms in the 2013 audiovisual data, 37 ms in the audiovisual 2015b data, and 16 ms in the 2015a audiovisual (flash-beep) data, and by 17 to 34 ms across the three speech experiments. The corresponding perceptual effects were significant in the mixed-effects models for AV-2013 (b = +0.39, *p* = 1.3×10), AV-2015b (b = +0.42, *p* = 4.4×10), speech E1 (b = +0.32, *p* = .003), speech E2 (b = +0.19, *p* = .034), and speech E3 (b = +0.33, *p* = 1.4×10), with each effect also supported by the participant-level test of the fitted ΔPSS (all *p* ≤ .036). In the 2015a dataset, the 16 ms shift was reliable in the per-participant test (*p* = .041) and in the source study’s own analysis, which we reproduce (F(1,9) = 10.8, *p* = .010), although its coefficient in the maximal mixed model fell just short of significance (b = +0.12, *p* = .080). In contrast, there was little evidence for a perceptual component when vision was paired with touch (a −2 ms shift, *p* = .30). For the audiotactile pairing, the perceptual shift was small (ΔPSS = 4 ms) and the two statistical approaches yielded different conclusions: the effect was significant in the mixed-effects model (*p* = .007), but not in the participant-level test (p = .48; see Table 2).

**Figure 4.**
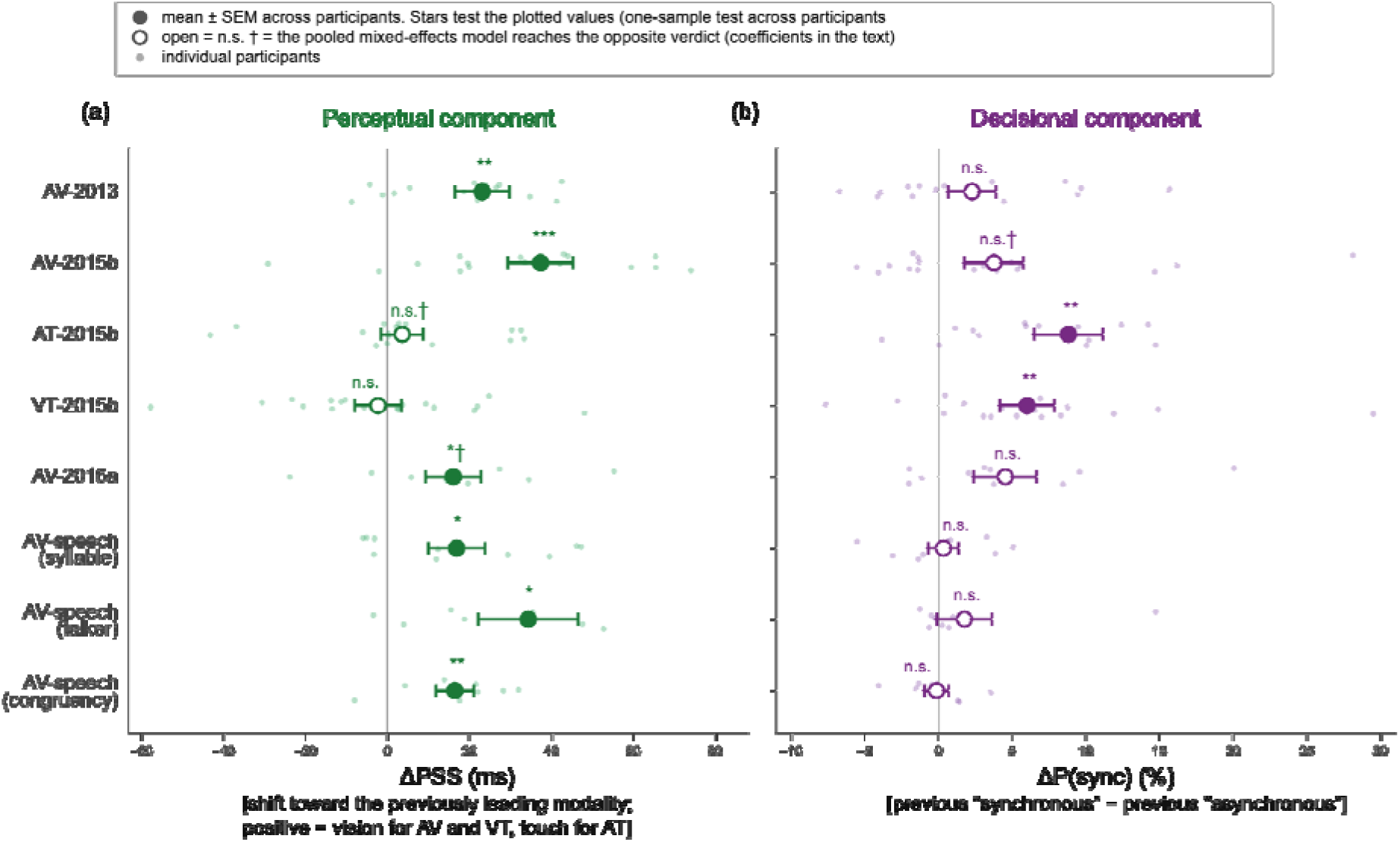
Magnitudes of the perceptual (left) and decisional (right) components across all eight datasets. For the perceptual component, the x-axis shows the change in point of subjective simultaneity (ΔPSS) as a function of the modality leading on the preceding trial, signed such that positive values indicate a shift toward the previousl leading modality (vision for audiovisual and visuotactile pairings; touch for audiotactile). For the decisional component, the x-axis shows the SOA-controlled change in the probability of a “synchronous” response (ΔP(sync)) following a previous “synchronous” versus “asynchronous” response. Small green and purple points show participant-level estimates for the perceptual and decisional components, respectively; larger symbols show th group mean ± between-participant SEM. The vertical reference line denotes zero effect. Asterisks give a one-sample test across participants of the plotted per-participant estimates (* p < .05, ** p < .01, *** p < .001), so the annotatio always agrees with the interval shown. Filled symbols indicate *p* < .05 and open symbols non-significant effects. † denotes a dataset in which the maximal mixed-effects model reaches the opposite verdict (the audiotactile and AV-2015a perceptual effects, and the AV-2015b decisional effect); the corresponding coefficients are given in the text.

**Table 2.**
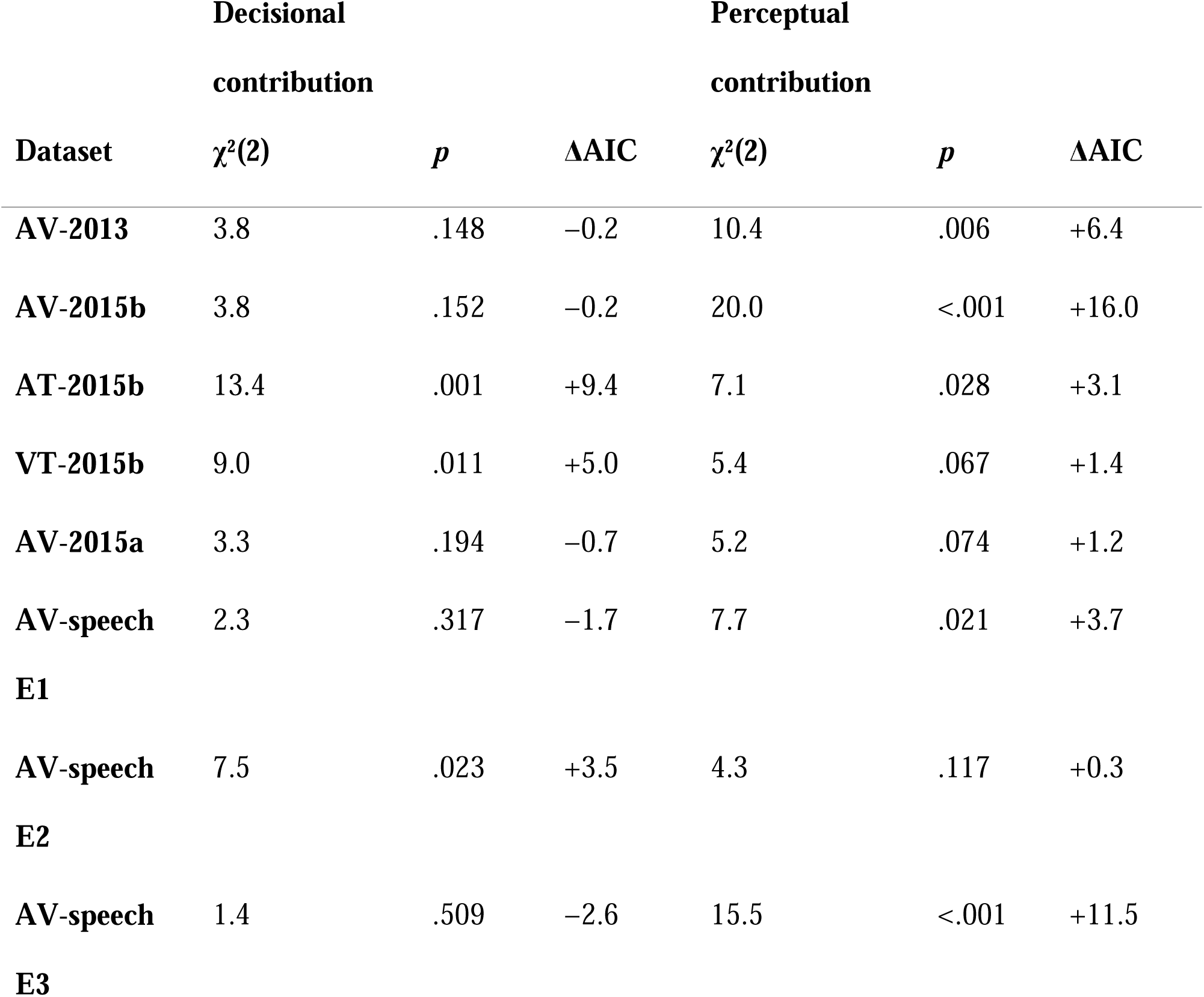
Unique contributions of the decisional and perceptual components to model fit across datasets. Each likelihood-ratio test compares the full model with a reduced model lacking the component indicated by the column heading, while retaining the same maximal random-effects structure. For the Decisional tests, the reduced model omits the previous-response terms. For the ‘Perceptual’ tests, the reduced model omits the previous-stimulus terms. Thus, a significant χ² indicates that the corresponding component explains variance not accounted for by the other component. χ² tests have 2 degrees of freedom. Positive ΔAIC values indicate better fit of the full model relative to the corresponding reduced model.

The decisional component showed the opposite profile. The previous response lifted the function by about 9% (mean P(sync) was 46% after a previous “synchronous” response and 37% after a previous “asynchronous” response, SOA-controlled) in the audiotactile data and 6% in the visuotactile data. Both effects were significant in the mixed-effects models (audiotactile b = +0.83, *p* = 1.8×10 ; visuotactile b = +0.57, *p* = .002) and were also supported by the participant-level analyses. For the audiotactile dataset, this conclusion was additionally confirmed by a penalised per-participant fit after the ordinary fit was distorted by response separation. In the audiovisual datasets, the decisional component was smaller, corresponding to 2 to 5%, and reached significance only in the 2015b audiovisual set and only in the mixed model (b = +0.47, *p* = .044; per-participant test p = .078), not in the AV-2013 or AV-2015a audiovisual sets (*p* = .33 and *p* = .12). In the speech experiments, the decisional main effect was near zero (all *p* > .39). As shown in the next section, however, these overall effects conceal a systematic dependence on stimulus content.

Together, the two components therefore trade off across the senses. The datasets with the weakest perceptual recalibration, the tactile pairings, carry the strongest decisional bias, and the datasets with strong perceptual recalibration carry little (Fig. 4).

### A trade-off between the senses, not between observers

Because the two components move in opposite directions across the senses, it is natural to ask whether the same trade-off is expressed at the level of individual observers. Do datasets showing stronger perceptual recalibration also show weaker decisional carryover, and, within datasets, do individuals who recalibrate more perceptually show less decisional carryover? The question can be asked at two levels, and the two levels give different answers.

At the level of the eight datasets, the two were moderately negatively related (r = −0.50; see Fig. 5): datasets with a larger perceptual component tended to show a smaller decisional component.

**Figure 5.**
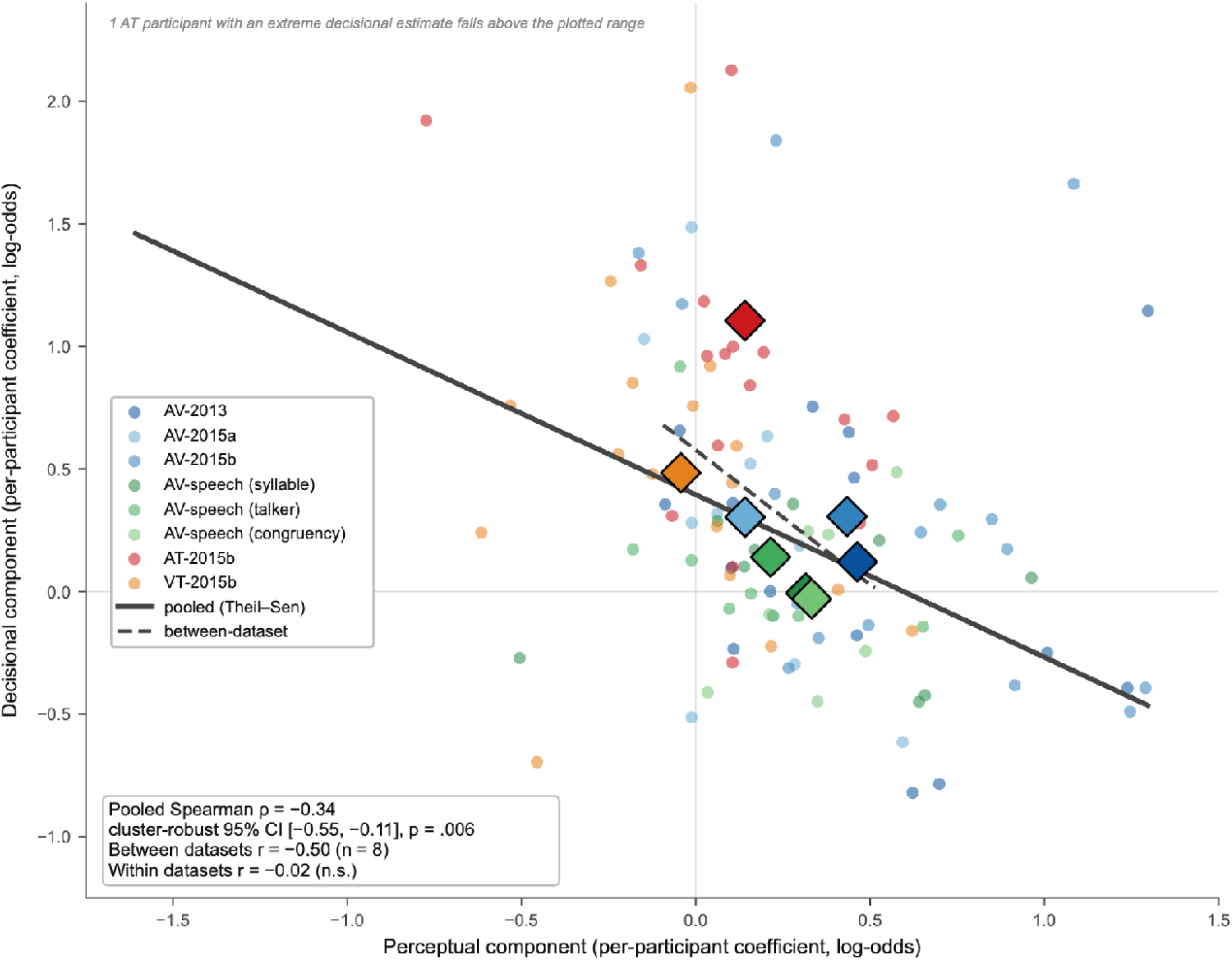
Participant-level relationship between the perceptual and decisional components. Each circle represents one participant within one dataset. The x-axis shows the participant-level stimulus-history coefficient indexing the odd, PSS-shift component; the y-axis shows the previous-response coefficient indexing the even, decisional carryover component. Both coefficients are expressed in log-odds and were estimated from penalised logistic models fitted separately to each participant’s trials. Colours identify datasets, and diamonds indicate dataset means. The solid line shows the robust Theil–Sen fit across all participant-by-dataset estimates; the dashed line shows the association across the eight dataset means (diamonds). The pooled association is negative (Spearman ρ = −0.34; participant-clustered 95% CI [−0.55, −0.11], *p* = .006), as is the association between dataset means (*r* = −0.50, *n* = 8), whereas the relationship disappears after removing between-dataset differences (*r* = −0.02, n.s.). Thus, the negative pooled relationship primarily reflects differences between sensory pairings rather than covariation between the two components across observers within a given dataset.

With only eight points, however, this does not reach significance (*p* = .20), but the direction is the clean mirror image of Fig. 4.

At the level of individual participants, the pattern is more informative. We fitted the same logistic model to each participant’s trials, yielding a perceptual and a decisional estimate for each dataset they completed. A few observers respond almost deterministically, repeating the previous verdict on nearly every trial, and for them an ordinary fit has no finite answer: the likelihood keeps improving as the coefficient runs off toward infinity. We therefore used Firth’s penalised logistic regression, which adds a small cost for extreme coefficients. This returns a usable estimate for every participant rather than forcing us to discard the most extreme responders, and it removes the bias that inflates logistic coefficients when each person contributes only a few hundred trials. The procedure yields 105 participant-by-dataset estimates from 53 distinct observers (fewer observers than estimates), because the same observers contribute to several datasets: the 2015b cross-modal study tested the same eighteen participants in all three pairings, and the three speech experiments drew on a shared pool of ten.

Two features of these estimates call for methods that do not assume a well-behaved scatter. Each rests on a single observer’s data and is therefore noisy, and a small number are extreme. We summarised the relationship in three ways, each insensitive to outliers in a different manner. The first is a Spearman’s rho rank correlation, which depends only on the ordering of the estimates and so cannot be dominated by one large value. Pooled across all estimates this yields ρ = −0.34. The second is a Theil–Sen line, which takes the slope between every pair of participants, 5,460 pairs in all, and uses the median of those slopes instead of minimising squared error. Because a median is unmoved by extreme values, almost a third of the points could be arbitrarily wrong without shifting the line, and nothing is assumed about the shape of the scatter. It gives a slope of −0.66 (95% CI −1.00 to −0.34). The contrast with ordinary least squares is instructive: that method returns a slope of only −0.20 and an ordinary correlation of r = −0.10, both flattened toward zero by a single observer with a very large decisional estimate. The robust measures agree with one another, the non-robust one understates the relationship.

The third analysis addresses dependence rather than outliers. The 105 estimates are not independent, since eighteen observers each appear three times, and a test that treated them as independent would call the relationship highly reliable. Resampling the 53 people rather than the 105 estimates gives an honest interval for Spearman’s ρ that is still negative and still excludes zero (95% CI −0.55 to −0.11, *p* = .006).

This pooled relationship is nevertheless inherited almost entirely from differences between the specific sensory combinations, not from differences between observers tested on the same pairing. Once each dataset is centred on its own mean, so that only within-dataset variation remains, the correlation vanishes (r = −0.02, *p* = .88), and the individual datasets’ correlations scatter around zero with no consistent sign. This is illustrated most clearly by the eighteen observers tested in all three cross-modal pairings: the same observers show predominantly perceptual recalibration for audiovisual stimuli, whereas decisional carryover is more prominent when the pairing involves touch. The trade-off is therefore associated with the sensory pairing rather than with stable individual differences between observers.

Across sensory pairings, stronger perceptual recalibration tends to go with weaker decisional carryover, and vice versa. Within individual datasets, however, the magnitudes of the two components are unrelated across participants. This pattern is consistent with the two components reflecting separable mechanisms with different sensitivities to sensory pairing, rather than alternative expressions of a common underlying individual tendency.

### Model evidence for separable perceptual and decisional contributions

A stronger test of the distinction between the two components is whether each explains variation in responses beyond that accounted for by the other. We tested this by removing one component at a time from the full model and asking whether the fit became reliably worse (Table 2). Removing the decisional component significantly worsened model fit for the two touch pairings, audiotactile (χ²(2) = 13.4, *p* = .001, ΔAIC = +9) and visuotactile (χ²(2) = 9.0, *p* = .011, ΔAIC = +5), and for one speech experiment. For the non-speech audiovisual datasets, removing the decisional component did not significantly reduce model fit (all *p*-values ≥ .15), indicating that the decisional terms provided little additional explanatory value beyond the perceptual terms. Removing the perceptual component showed the complementary pattern: it reliably worsened the fit for the AV-2013 and AV-2015b datasets and for two of the three speech experiments (all *p*- values ≤ .021), and fell marginally short for AV-2015a (*p* = .074) and visuotactile (*p* = .068). Audiotactile was the single dataset in which both components earned their place, marking it as genuinely mixed. Across the eight datasets, the perceptual component improved the fit in five and the decisional in three, so the perceptual effect is the broadly robust one and the decisional is real but concentrated in the touch pairings, exactly where the summary in Fig. 4 places it. This is the strongest form of the claim, because it shows that where the decisional component is significant it is not simply a repackaging of the perceptual effect.

### The AV-speech decisional carryover (E1) is tied to content, not to identity, and lasts a single trial

If the decisional carryover is genuinely a decision being carried forward, rather than a habit of repeating the previous response, it should depend on what is being judged. The speech experiments test this, because successive trials could repeat or change the spoken syllable (E1; Fig. 6) and, in a separate experiment, the identity of the talker (E2; statistics below, not shown in Fig. 6). The decisional carryover was tied to the message, not the messenger. When the syllable repeated, the previous verdict carried over (Fig. 6a); when the syllable changed, the carryover collapsed (Fig. 6b). This gating by syllable is a large effect and it survives the conservative per- participant test (interaction b = +0.78, t(9) = 2.5, *p* = .032). The ΔP(sync) values shown within the individual panels of Fig. 6 are each condition’s carryover considered alone, and none of these subset-level simple effects is individually reliable; the gating claim rests on the interaction, the within-participant contrast between the same- and different-syllable conditions, marked by the bracket in Fig. 6. Changing the talker while leaving the syllable’s role intact did not reliably gate the carryover (interaction b = +0.27, *p* = .23): the verdict transferred across a change of voice. The congruency experiment (E3), which manipulated neither in this way, showed no reliable carryover to gate (*p* = .51).

**Figure 6.**
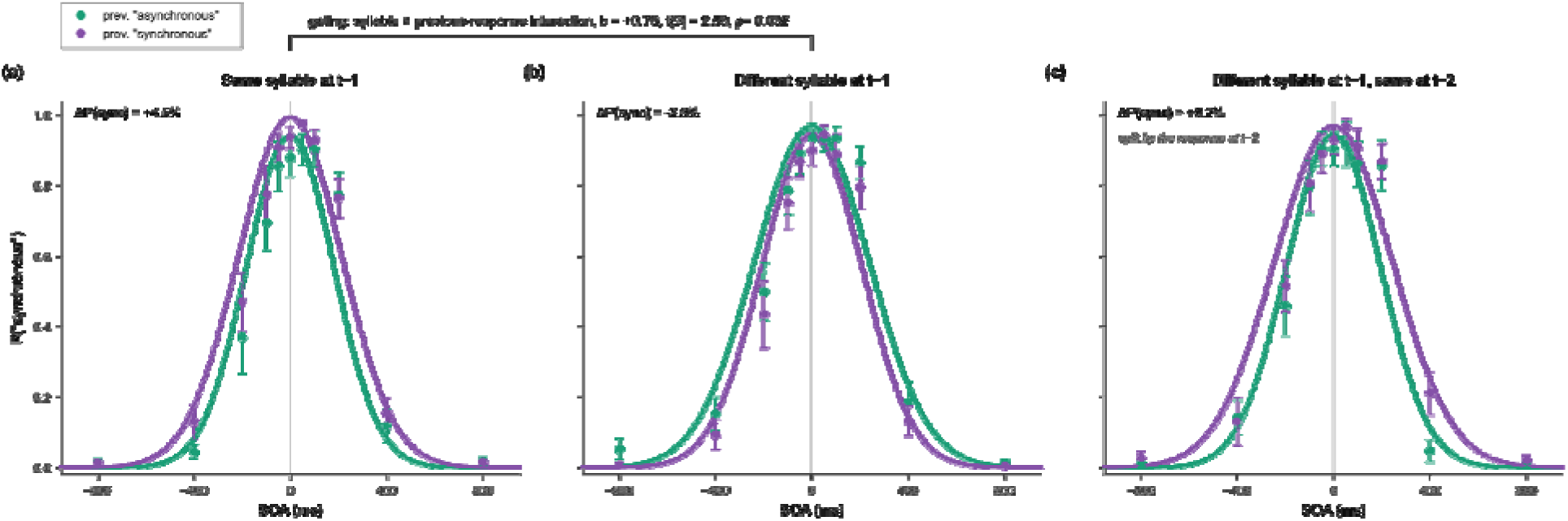
Content specificity and temporal extent of decisional carryover in audiovisual speech (E1). The x-axis shows current-trial stimulus onset asynchrony (SOA ms), and the y-axis gives the probability of a “synchronous” response. Green and purple symbols and curves show trials conditioned on a preceding “asynchronous” versus “synchronous” response, respectively (in (c), the response two trials back). In (a), the current and immediately preceding trials contained the same syllable; the vertical separation between the curves shows the decisional carryover. In **(b)**, the syllable changed from the preceding trial, and this separation was absent. In **(c)**, analyses were restricted to trials for which the immediately preceding syllable differed from the current syllable but the syllabl two trials back matched; here, the curves are conditioned on the response two trials back and show no recovery of the decisional carryover across the intervening non-matching trial. Symbols show group mean response probabilities at each SOA ± between-participant SEM, and solid curves show the corresponding fitted simultaneity functions. The ΔP(sync) value in each panel is that condition’s SOA-controlled carryover (in %); the bracket linking panels (a) an (b) reports the syllable × previous-response interaction, the test of the gating itself.

We also asked whether a verdict can reach across an intervening trial, re-emerging when a syllable returns two trials later even after a different syllable has come between. It does not.

Restricting to trials on which the previous syllable was a switch, the verdict from two trials back produced no carryover when its syllable matched the current one, and matching made no difference (interaction b = +0.37, *p* = .17). The same was true for the talker in the talker experiment (two-back interaction *p* = .71). The decisional carryover is therefore a one-step, content-specific effect: it depends only on the immediately preceding trial, and only when the judgment-relevant content, the syllable, repeats.

## Discussion

Decomposing serial dependence in a simultaneity judgment into a perceptual component, a sideways shift of the PSS, and a decisional component, a vertical bias to repeat the last verdict, reveals structure that the raw PSS conceals. The two components show distinct modality profiles: perceptual recalibration is most consistently observed for audiovisual stimuli and is weaker for pairings involving touch (see also Alais et al. (2017a)), whereas the decisional carryover is confined to the tactile pairings and is unreliable for audiovisual pairs. Across datasets the two trade off, although this is a property of the modality rather than of the individual. In speech, the decisional carryover is gated by the spoken syllable but not by the talker identity, and we found no evidence that it extends across an intervening trial, marking it as a content-specific, one-step decision memory rather than a motor repetition bias or a persistent multi-trial influence.

The central result of this study is that the shift in PSS, typically treated as a unitary measure of temporal recalibration, comprises at least two separable components, and that separating them re- draws the map of temporal recalibration. When the components are partialled, perceptual recalibration and decisional carryover do not merely differ in size; they are selectively associated with different sensory pairings. This is difficult to see without the decomposition, because the undecomposed PSS sums a perceptual shift that is large for audiovisual timing with a decisional bias that is large for touch, blurring two opposite gradients into one.

Particularly strong evidence that the two components capture separable contributions comes from the case where they coincide. Asking of each dataset whether a component explains variance the other cannot, audiotactile timing is the single case in which both components earn their place, marking it as genuinely mixed. This matters because a single underlying process, viewed two ways, is unlikely to require both terms at once: wherever one description sufficed, the other would be redundant. That the model needs both for audiotactile timing, while needing only the perceptual term for audiovisual pairings and only the decisional term for visuotactile, is what a genuine dissociation looks like when the two processes happen to overlap. The audiotactile perceptual contribution is nonetheless small, a few milliseconds, and reaches significance in the mixed model without being confirmed by the more conservative per-participant test, so audiotactile timing is best read as a pairing in which a weak perceptual realignment coexists with a strong decisional bias, rather than one in which both are substantial.

The predominance of the perceptual component in audiovisual timing is also consistent with earlier experimental evidence that rapid audiovisual recalibration does not require a judgment on the preceding trial. Van der Burg et al. (2013) observed recalibration even when no temporal judgment was made on that (passive) trial, and subsequent work showed that recalibration followed the physical temporal order even when observers judged that order incorrectly (Van der Burg et al., 2018). Thus, audiovisual recalibration appears to depend primarily on the preceding stimulus rather than on the observer’s decision about it. This is notable because serial dependence in other domains can depend strongly on the task performed on the preceding trial, including judgments of facial attractiveness (Van der Burg et al., 2019), visual motion (Alais et al., 2017b), and prosody (Van der Burg et al., 2024). The relative independence of audiovisual recalibration from the preceding judgment therefore fits well with the present finding that audiovisual timing is dominated by the perceptual (odd) component, with comparatively little decisional (even) carryover.

Why perception should realign audiovisual timing readily but tactile timing hardly at all, while the decisional bias runs the other way, remains an open question, but the pattern suggests a functional account. Audiovisual asynchronies are pervasive in the natural world, because sound arrives later than light from any distant source, making continual adjustment of their relative timing potentially useful. Where the perceptual system does not re-align the senses, the observer may lean instead on a simpler strategy, repeating the judgment reached a moment earlier. According to this interpretation, the cross-modal trade-off reflects a division of labour between perceptual and decisional mechanisms. This interpretation remains provisional, however, given that the cross-modal comparison is based on only eight datasets.

The syllable gating results (AV-speech E1, Fig. 6) provide insight into the representational basis of the decisional carryover. A purely motor account, in which observers simply tend to repeat the previous key press, would predict carryover irrespective of the stimulus being judged. Instead, carryover depended on whether the spoken syllable repeated, indicating that it was tied to judgment-relevant stimulus content. It is not a generic sense of familiarity from any repeated feature, because repeating the talker did not modulate carryover. And it is not a multi-trial episodic store, because a verdict does not reach across an intervening different-syllable trial. What remains is a one-step, content-addressed decision memory. Observers carry forward the verdict reached about this content on the immediately preceding trial and apply it again when the same syllable recurs.

That the carryover follows the syllable and ignores the talker is itself informative. The syllable is the thing whose audiovisual timing is being judged; the talker is incidental to that timing, because the same syllable has much the same lip-to-sound relationship whoever speaks it. The decision memory is thus indexed to the feature that carries the timing information and generalises over the feature that does not, which places it at a talker-invariant, phonetically specific level of representation, the level at which speech perception is known to normalise across voices (Miller, 1989, Choi et al., 2018, Kraljic et al., 2008). Because the original design varied syllable and talker but did not fully cross them, the cleanest test of this message-versus- messenger gating would come from a design that crosses the two factors, which we suggest as a direct next step.

Throughout, we have treated the sideways shift of the PSS as the perceptual component and the vertical bias as the decisional one, and it is worth being explicit that this labelling concerns geometry, not mechanism. A PSS shift can itself arise from a decision process rather than a change in perceived timing, as criterion-based accounts of temporal-order and simultaneity judgments have argued: an observer who revises which asynchronies count as synchronous will move the PSS with no change in experience (Yarrow et al., 2011, Yarrow et al., 2015). Notably, rapid audiovisual recalibration follows the physical rather than the perceived temporal order of the preceding trial (Van der Burg et al., 2018), so the realignment does not require awareness of the preceding asynchrony, which sits more naturally with a stimulus-driven realignment than with a deliberate revision of the decision criterion.(Yarrow et al., 2011, Yarrow et al., 2015) Our decomposition does not by itself settle whether the odd component is perceptual in that deeper sense. What it adds is the isolation of a second, logically separate source of serial dependence, the even repetition bias, which standard PSS-based analyses miss entirely and which is decisional in a different and unambiguous way: it is a carried-forward verdict, not a re-drawn boundary. The two senses of "decisional" should therefore be distinguished, because the present decomposition separates a carried-forward response tendency from a shift in the decision criterion.

While the findings are consistent across complementary analyses, some of the finer-grained claims require qualification. The core dissociation between perceptual and decisional carryover is evident across complementary analyses and does not depend on any single method of estimation. Greater caution is warranted, however, when interpreting the magnitude and generality of these effects. Participant numbers vary across datasets, with ten or fewer observers in AV-2015a and the speech experiments, limiting the sensitivity of the between-participant analyses, while the perceptual and decisional effect magnitudes derived from Gaussian fits are necessarily approximate. Two effects sit close enough to the threshold that they should not be read as established: the audiotactile perceptual shift and the audiovisual decisional lift in the (Van der Burg et al., 2015b) dataset each reach significance in the mixed model but not in the more conservative per-participant tests. The proposed cross-modal trade-off is more tentative still: it is based on only eight dataset-level observations and is not itself statistically reliable. Likewise, the speech results strongly suggest content-specific gating, but the original design did not fully cross syllable and talker identity, limiting the precision with which the underlying representation can be identified. These considerations leave the central perceptual–decisional dissociation intact, while placing appropriate limits on the stronger claims concerning the magnitude, cross-modal organisation, and representational specificity of its component processes.

## Conclusion

A simultaneity judgment carries at least two kinds of memory of its recent past: a stimulus- linked (perceptual) shift in the estimated point of simultaneity and a decisional tendency to repeat the last verdict. Separating these components within the GLMM shows that they are not redundant measures of a single effect, but dissociable processes with distinct modality profiles and different dependencies on recent stimulus history: dissociable enough that audiotactile timing requires both at once. The decisional carryover in particular emerges as a content- specific, one-step decision memory that can be distinguished from perceptual temporal realignment. Applying this decomposition to other existing datasets will be important for establishing how much of what has previously been described as rapid temporal recalibration reflects perceptual realignment, decisional carryover, or some combination of the two. Distinguishing these contributions should provide a more precise account of how recent experience shapes cross-modal timing judgments across sensory pairings and experimental contexts.

## Acknowledgements

We thank the co-authors of the source studies for making their data available.

## Author Contributions

J.C.: conceptualisation, methodology, formal analysis, visualisation, writing – original draft. E.V.d.B.: investigation (collection of the original datasets), data curation, writing – review and editing.

## Ethics Statement

This article reports a secondary analysis of previously published data (see Table 1). No new data were collected. Each original experiment was approved by the human research ethics committee of the institution at which it was conducted, and all participants gave informed consent, as reported in the source articles.

## Data Availability

The trial-level data for all eight datasets and the analysis code (R and Python) are available at https://osf.io/9zcfj/overview?view_only=9cb51621265f4d9f91b4033f7f63b964

